# Impairment of Neuronal Activity in the Dorsolateral Prefrontal Cortex Occurs Early in Parkinsonism

**DOI:** 10.1101/2024.10.22.619684

**Authors:** Noah Hjelle, Biswaranjan Mohanty, Tanner Hubbard, Matthew D Johnson, Jing Wang, Luke A Johnson, Jerrold L Vitek

**Affiliations:** Department of Neurology, University of Minnesota, Minneapolis, MN, USA; Department of Biomedical Engineering, University of Minnesota, Minneapolis, MN, USA

## Abstract

**Background:** Parkinson’s disease (PD) is often characterized by altered rates and patterns of neuronal activity in the sensorimotor regions of the basal ganglia thalamocortical network. Little is known, however, regarding how neuronal activity in the executive control network of the brain changes in the parkinsonian condition.

**Objective:** Investigate the impact of parkinsonism on neuronal activity in the dorsolateral prefrontal cortex (DLPFC), a key region in executive control, during a go/nogo reaching task.

**Methods:** Using a within-subject design, single and multi-unit neuronal activity was recorded in the DLPFC of a nonhuman primate before and after the induction of mild parkinsonism using the neurotoxin 1-methyl-4-phenyl-1,2,3,6-tetrahydropyridine (MPTP).

**Results:** Coincident with development of mild parkinsonian motor signs, there was a marked reduction in the percentage of DLPFC cells with significant task-related firing rate modulation during go and nogo conditions.

**Conclusions:** These results suggest that DLPFC dysfunction may occur early in parkinsonism and contribute to cognitive impairments and disrupted executive function often observed in PD patients.

## Introduction

Parkinson’s disease (PD) is a neurodegenerative disorder characterized by disruptions in motor function, e.g., delayed movement initiation, decreased movement speed, rest tremor, and increased joint rigidity^1^. Non-motor symptoms, however, such as cognitive dysfunction, are also prevalent components of PD^2,3^. These cognitive impairments include changes in executive functions such as working memory, set shifting, and movement inhibition^4–6^. Functional imaging studies have shown that the DLPFC, a critical node in the BGTC network involved in executive function, is impaired in PD^5,7–10^. This impairment is likely secondary to the role of dopamine in mediating executive functions involving the DLPFC^11,12^. While some functional imaging studies suggest parkinsonism results in hypoactivation in the prefrontal cortex^13,14^, consistent with classic models of basal ganglia function in which excessive inhibitory activity from the internal segment of the globus pallidus is hypothesized to result in reduced excitatory thalamo-cortical drive^15,16^, other studies have found hyperactivation in DLPFC in PD patients compared to controls^5,17,18^. There are limited neuronal data at the single unit level characterizing changes in DLPFC activity in PD to support or refute either of these findings.

One task known to probe executive function in the context of DLPFC is the go/nogo task, which require subjects to discern between two target types that indicate either taking or avoiding an action^19,20^. Additionally, the go/nogo paradigm has been useful to show alterations in executive function in PD, such as movement preparation and response inhibition^19,20^. In this study, a go/nogo touch screen task was used to engage DLPFC, and to test the hypothesis that neuronal processing in the DLPFC is abnormal in early PD through the induction of a mild parkinsonian state. We compared task-related single and multi-unit neuronal firing characteristics in the DLPFC of a nonhuman primate before and after induction of mild parkinsonism using the neurotoxin 1-methyl-4-phenyl-1,2,3,6-tetrahydropyridine (MPTP).

## Methods

### Surgical procedures

All procedures were approved by the University of Minnesota Institutional Animal Care and Use Committee and complied with US Public Health Service policy on the humane care and use of laboratory animals. One adult female rhesus macaque (Macaca mulatta, 20 years of age) was used in this study. Surgery was performed under isoflurane anesthesia using aseptic techniques. The animal was implanted in the right DLPFC with a 96-channel Utah microelectrode array (Pt-Ir, 1.5 mm depth, 400 um inter-electrode spacing, Blackrock Microsystems) using surgical methods described previously^21–23^. DLPFC was identified based on sulcal landmarks during the array implantation surgery (Fig. 2A).

### Go/nogo task and data collection

The animal was trained to perform a visually cued go/nogo reaching task (Fig 1A). Trials were initiated when the animal placed its left hand on a capacitive touchpad (“start-pad”) and, after a two second delay, a cue appeared in one of three randomly selected locations. Two seconds following this cue a “go” target appeared at the selected location in 80% of the trials, and a “nogo” target would appear in 20% of the trials. A successful go trial required the primate to leave the start-pad within 1.5 seconds and touch the target within another 1.5 seconds. A successful nogo trial required the animal to hold on the start-pad for 1.5 seconds following target presentation. Successful trials resulted in a juice reward. Reaction time was defined as the time between presentation of the go target and reach initiation (the time when the animal’s hand left the start-pad). Reach duration was defined as the time between reach initiation and contact with the target. The animal initiated the next trial by voluntarily returning to the start-pad. Go and nogo target appearance timepoints were extracted from the task software and reach initiation timepoints were recorded from the start-pad. Raw neurophysiological data were collected using a TDT workstation (Tucker Davis Technologies) operating at ∼ 25 kHz sampling rate. Activity of DLPFC units was recorded while the animal was seated, head fixed, in a primate chair performing the reaching task.

**Fig 1.**
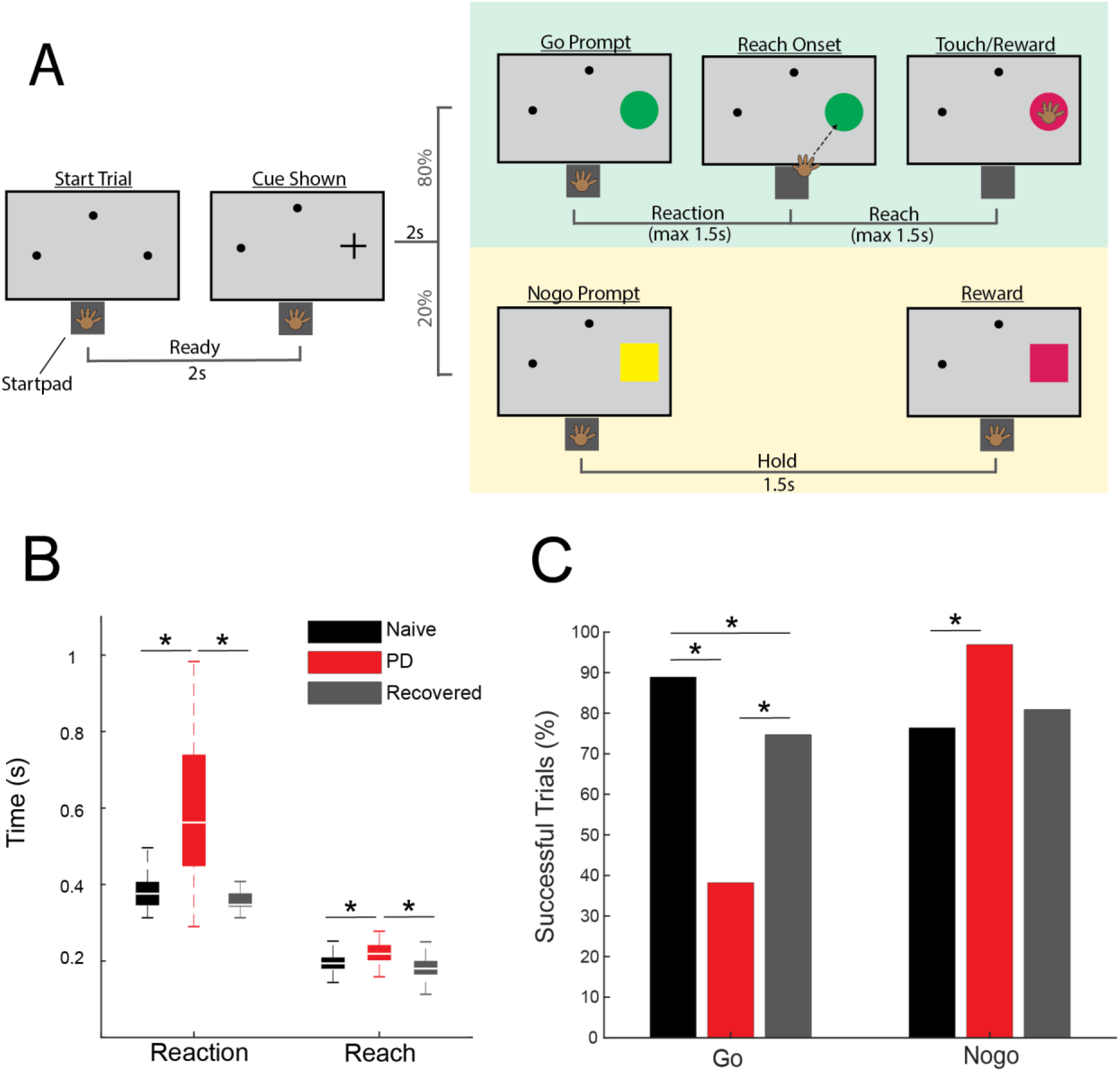
**(A)** Go/nogo task paradigm. **(B)** Reach and reaction times during successful go trials in naive (black), PD (red), and recovered (gray) (WRS, p < 0.05). **(C)** Percentage of successful go and nogo trials (Χ^2^, p < 0.05).

Once data were collected in the normal state, the animal was rendered parkinsonian by one intracarotid injection of MPTP (0.4mg/kg). Overall parkinsonian severity was assessed using a modified Unified Parkinson’s Disease Rating Scale (mUPDRS), which rated appendicular motor symptoms (upper and lower limb rigidity, bradykinesia, akinesia, and tremor) on the hemi-body contralateral to neural recordings using a 0-3 scale (0 = normal, 3 = severe, maximum total score = 27)^24^. We observed mild parkinsonian signs (mUPDRS: 2.8±1.14, 5 ratings) after MPTP injection; however motor signs improved over subsequent weeks and returned to a baseline mUPDRS score of 0 after 11 days (defined here as the “recovered” state). Post-MPTP neural data were obtained in both the mild parkinsonian and recovered states.

### Statistical analysis of neuronal data

Neuronal recordings were analyzed offline using custom software developed in MATLAB (Mathworks) and Offline Sorter (Plexon). Raw data were bandpass filtered 300-5000 Hz, and single and multi-units were isolated and sorted using principal component and template-based methods in Offline Sorter (hereafter referred to collectively as “units”). Spike trains were aligned to go target appearance, nogo target appearance, or reach onset. A trial-averaged spike density function for each unit was generated by convolving each spike with a gaussian kernel (60 ms variance) and a time resolution of 10 ms. The baseline firing rate for each unit was defined as the mean of the spike density function during a 1.5 second period beginning 0.25 seconds after trial initiation, before cue presentation. Units with an extremely low firing rate were excluded from further analysis (less than 0.75 spikes per second). To investigate neuronal modulation during the reaction time period, spiking activity immediately following target appearance until the minimum reaction time across all trials (255 ms) was used. The same time frame was used for nogo trials. To investigate neuronal modulation during reach initiation, activity 50 ms prior to reach onset until the minimum reach duration across all trials (144 ms) was used. A one-sample 2-tailed t-test was used to compare the mean baseline firing rate of each unit to the firing rates during the reaction or reach periods. Units with a significant change in firing rate during the analysis window (p ≤ 0.002) compared to baseline were classified as modulated.^25^ With a time resolution of 10 ms and a maximum window of 255 ms, the maximum number of comparisons was 25, so 0.002 (0.05/25) was chosen as a conservative threshold for determining whether a cell was modulated. Units with a significant increase in firing rate were further classified as activated, and those with a significant decrease as suppressed. Reaction times were compared across states using the Wilcoxon rank sum test (WRS), as were the reach durations. Chi-squared tests [Χ^2^(DoF, N)] were used to compare the percentage changes in the number of modulated, activated and suppressed units, the ratio of activated over suppressed units, and changes in task success rates, between the naïve, PD, and recovered conditions.

## Results

### Effects of MPTP on task performance

MPTP administration induced a mild parkinsonian state based on clinical assessments (2.8±1.14, 5 ratings). In the recovered state the mUPDRS score returned to zero. The NHP completed 288 go trials and 72 nogo trials in the naïve state and recovered state. In the parkinsonian condition the NHP completed 123 go trials and 31 nogo trials. As indicated in Fig. 1B, reaction times increased in the parkinsonian condition compared to the naïve (WRS; z = -8.2862, p < 0.001) and recovered states (WRS; z = 8.7851, p < 0.001). Reach times were also longer in the parkinsonian condition compared to naïve (WRS; z = -5.4332, p < 0.001) and recovered states (WRS; z = 6.1389, p < 0.001). There was a decrease in task success rate during go trials from 81% in naïve to 36% in the parkinsonian condition (Fig. 1C, *left*) [Χ2(1,411) = 85.1, p < 0.001]. Task success rate during go trials was higher in the recovered state compared to the parkinsonian condition [Χ^2^(1,411) = 38.0233, p < 0.001]. Nogo trial success rates were increased in the parkinsonian condition compared to the naïve state [Χ^2^(1,103) = 6.2442, p = 0.0125]. In the recovered state, nogo task performance returned to a level similar to that observed in the naïve state (Fig. 1C, *right*).

### Parkinsonism alters neuronal modulation in DLPFC

A total of 410 units in DLPFC were recorded in this study (n = 149 naïve, n = 101 parkinsonian, and n = 160 recovered. Representative neurons (Fig. 2B) illustrate our main finding that there is a significant reduction in task-related unit activity in DLPFC in the mild parkinsonian condition. While 62.4% of units had significant firing rate modulation during the go reaction time period in the naïve state, in the parkinsonian condition only 24.8% were modulated ([Χ^2^(1,250) = 34.2638, p < 0.001]). Similarly, there was a reduction in the percent of units with significant modulation in the go reach period (53.7% in naïve compared to 31.7% in the parkinsonian condition, [Χ^2^(1, 250) = 11.7901, p < .001]) (Fig. 2C). During the recovered state the percent of units with a significant firing rate modulation returned to levels similar to the naïve state for both go reaction (naive: 62.4%, PD: 24.8%, recovered: 61.9%) [Χ^2^(1,261) = 34.2148, p < 0.001] and go reach periods (naive: 53.7%, PD: 31.7%, recovered: 49.4%) [Χ^2^(1,261) = 7.9289, p = 0.005]. Modulation during the nogo reaction period decreased from 29.7% in the naïve state to 12.0% in the parkinsonian condition [Χ^2^(1, 250) = 10.7307, p = 0.001], but did not return to naïve levels in the recovered state [Χ^2^(1,261) = 0.9288, p = 0.3352].

**Fig 2.**
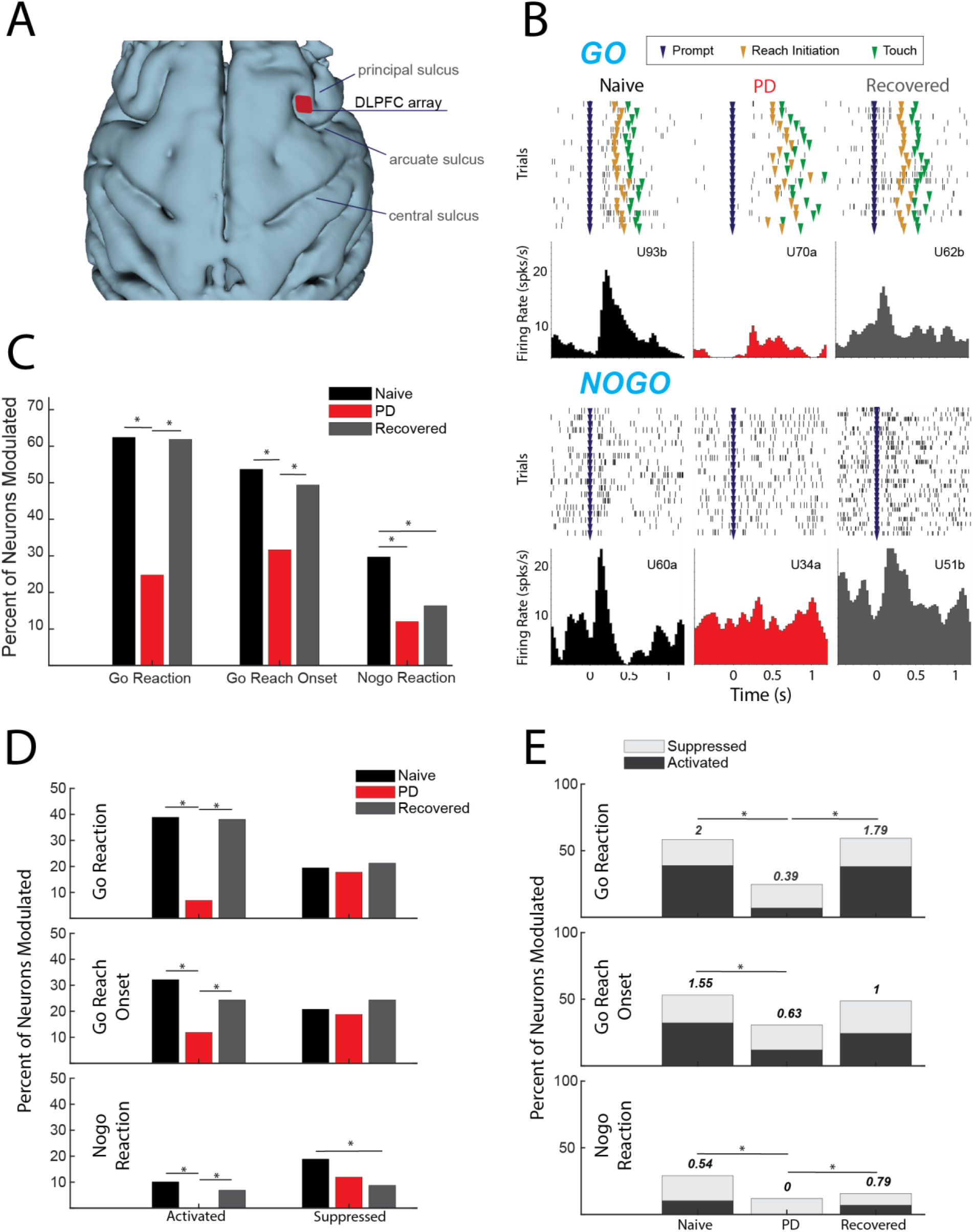
**(A)** 3D reconstruction of cortex with DLPFC array location. **(B)** Example cells during go (*upper*) and nogo (lower) trials from each condition. **(C)** Percentage of cells modulating during comparison window (p < 0.05, Χ^2^). **(D)** Percent of total cells activated (left) and suppressed (right) (p < 0.05, Χ^2^). **(E)** Ratio of percent activated over percent suppressed, indicated by number above the bars (p < 0.05, Χ^2^).

### Parkinsonism decreases neuronal activation

The results presented in Figure 2C showed that fewer DLPFC cells were modulated in the mild PD condition, irrespective of whether that modulation was due to significant increases (activation) or decreases (suppression) in firing rate. We then examined how the parkinsonian state impacted the proportion of cells classified into these modulation subcategories (Fig. 2D) As described below, we found that the reduction in modulation in the PD condition was driven predominantly by a loss of cells that were significantly activated during the go/nogo task.

### Activation

The percentage of cells activated during the go reaction period decreased from 38.9% in the naïve state to 6.9% in the parkinsonian condition [Χ^2^(1,250) = 32.0287, p < 0.001)] and the percentage of cells activated during the reach period decreased from 32.2% to 11.9% [Χ^2^(1,250) = 10.0132, p < 0.001]. In addition, the percentage of cells activated during the nogo reaction period decreased to zero from naïve to the parkinsonian condition [Χ^2^(1,250) = 10.7876, p < 0.001]. In the recovered state the percentage of activated units increased back to naïve levels during go reaction (38.1%) [Χ^2^(1,261) = 31.2728, p < 0.001], reach (24.4%) [Χ^2^(1,261) = 6.1473, p = 0.0132], and nogo reaction (6.9) [Χ^2^(1,261) = 7.2251, p = 0.0072].

### Suppression

There was no significant difference in the percentage of suppressed units during the go reaction period [Χ^2^(1,250) = 0.1062, p = 0.7445], reach [Χ^2^(1,250) = 0.1495, p = 0.699], or the nogo reaction period [Χ^2^(1,250) = 2.1119, p = 0.1462] between naïve and the parkinsonian condition. Similarly, there was no significant difference in the percentage of suppressed units between the parkinsonian condition and recovered state during the go reaction period [Χ^2^(1,261) = 0.4561, p = 0.4994], reach period [Χ^2^(1,261) = 1.1087, p = 0.2924], or nogo reaction period [Χ^2^(1,261) = 0.6939, p = 0.4048]. During the nogo reaction period there was a significant decrease in suppression between the naïve and recovered states [Χ^2^(1,309) = 6.6395, p = 0.01] (Fig. 2D).

### Ratio of activation to suppression

The loss of activation without a significant change in suppression resulted in a decrease in the ratio of activation to suppression from naïve to the parkinsonian state during all three analysis periods (Fig. 2E). This ratio decreased from 2 to 0.39 during go reaction [Χ^2^(1,118) = 13.6973, p < 0.001], 1.55 to 0.63 during reach [Χ^2^(1,112) = 6.7274, p = 0.01], and 0.54 to 0 during nogo reaction [Χ^2^(1,56) = 6.1091, p = 0.0134]. In the recovered state, the ratio of activated to suppressed cells increased during go reaction [Χ^2^(1,124) = 11.6236, p < 0.001] and nogo reaction [Χ^2^(1,38) = 8.0947, p = 0.004], returning closer to that observed in the naïve state.

## Discussion

The present study investigated the effects of MPTP-induced parkinsonism on neuronal activity in the DLPFC of a nonhuman primate. A unique advantage of this animal model not feasible in human studies is that it allows for a within-subject comparison of changes in activity of neuronal cell populations between healthy and parkinsonian conditions. Induction of the parkinsonian state was associated with a decrease in task-dependent neuronal modulation of firing rates. This reduced modulation was driven by a decrease in the number of activated neurons, leading to a decrease in the ratio of activated to suppressed neurons. The recovered state was associated with an increase in task-dependent modulation, driven by increased activation, compared to the parkinsonian condition. Importantly, these changes in neural activity occurred even in a mild state of parkinsonism, suggesting that the mechanisms involved in cognitive dysfunction may be initiated in the early stages of PD.

### Comparison to previous studies of DLPFC in PD

Our results suggest DLPFC is hypoactive during motor preparation and execution in mild parkinsonism. Consistent with this observation, studies utilizing PET and fMRI have found decreased activation during both self-initiated and externally cued timing tasks in the right DLPFC in PD patients compared to healthy control subjects^13,14,26^. These groups hypothesized that the decrease in DLPFC activation is a result of reduced thalamic output to the cortex as a result of decreased dopamine in the basal ganglia, consistent with the classical model of PD pathophysiology^15^. Nevertheless, there are multiple imaging studies that have identified increased rather than decreased activation in the DLPFC during motor tasks^5,17,18^. Martin et al. used fMRI to identify increased activation in the DLPFC during motor planning in early-stage PD patients performing a finger tapping task, but found no change in DLPFC activity during movement execution^17^. Similarly, Disbrow et al. found increased BOLD signal in DLPFC bilaterally prior to un-cued movement^5^. Some suggest that this relative hyperactivity in DLPFC could be a compensatory mechanism to accommodate for disrupted function of motor areas in PD^17,18^. Evidence of compensatory mechanisms in these studies were supported by a lack of change in task performance despite increased motor symptoms based on UPDRS-III motor signs, though this was not a phenomenon observed in the present study (i.e., motor signs were observed and quantified by clinical exam as well as during the task).

While our findings are consistent with the classical model of PD, it is also possible that though our task clearly engaged DLPFC, it may not have required the conceptualization of movement prior to target appearance that may involve compensatory mechanisms, as suggested by Martin et al.^17^. Furthermore, the high success rate in the parkinsonian condition suggests that the primate may not have been habituated to the go condition, and therefore may have been simply waiting for the go cue to appear before planning movement^27^. There is also the possibility that the parkinsonian condition obtained was too mild to have induced any compensatory effects. Some studies, however, finding hyperactivation in DLPFC suggested compensatory effects were present in mild PD patients^17,18^. While we did not find evidence of an overactive DLPFC in the PD state as might be hypothesized based on these imaging studies, future studies are necessary to fully probe the hypothesis of compensatory mechanisms triggered in the frontal cortex in PD. For example, while our data was collected in a mild PD state, more severe PD states should be investigated to determine whether hyperactivity develops when motor signs are more severe, and compensation is necessary to perform the task.

### Recovery from MPTP injection

Recovery from a mild state of parkinsonism following MPTP administration has been previously documented^28^. The mechanisms of this recovery, however, are not fully understood. Hypotheses include reactive synaptogenesis (temporary, quick onset synapse formation) and denervation hypersensitivity (increased sensitivity to a neurotransmitter after loss of synapses), and uptake of excess dopamine in the nigrostriatal circuit^29–31^. The most likely mechanism underlying this recovery however, is that MPTP administration causes cell injury, but some cells recover over time^31^. By including the recovered condition in this study we were able to show a possible correlation between DLPFC activity and go/nogo task behaviors such as success rate and reaction time. Although the changes in DLPFC activity in the parkinsonian condition that resolved following recovery provide compelling evidence is support of the role of DLPFC deficits in the observed motor dysfunction, whether the change in behavior was the cause or the result of the change in DLPFC activity needs to be determined with additional studies.

### Limitations and future directions

This study included only one NHP, but represents our early findings that are part of a larger study where multiple animals are being enrolled to validate these findings. Another potential limitation is that the task may not have probed the response inhibition aspects of the DLPFC that we had intended, which may be reflected in the increased success rate of nogo trials in the parkinsonian condition. In the future we will modify the task paradigm to induce response inhibition while allowing us to investigate the changes in the DLPFC in early PD where the animals are still able to perform the task. There are challenges, however, to designing tasks that are both cognitively complex and feasible for a parkinsonian animal given their impaired cognitive and motor functions. Regardless, this study provided data to support the finding that even in a mild disease state there are salient changes to neural activity in the DLPFC. In the future we may look to quantify cognitive performance of the primate while parkinsonian to characterize neurophysiological changes related specifically to cognitive disruption and identify how dopamine replacement therapy and DBS alter pre-frontal cortical activity. While deep brain stimulation is effective at modulating motor cortex activity in PD patients, its effect on frontal cortical regions is less well understood^32^. Understanding the neural mechanisms underlying frontal cortical dysfunction in PD will help motivate and inform neuromodulation techniques that would allow us to improve neural function for both parkinsonian motor and cognitive behaviors.

## Acknowledgement

We would like to thank our colleagues in the Neuromodulation Research Center for helpful comments and critiques related to this study and especially thank our animal core team of Claudia Hendrix, Hannah Baker, Adele DeNicola, and Elizabeth McDuell as well as our veterinary and animal care colleagues at the University of Minnesota Research Animal Resources (RAR).

## Ethical Compliance Statement

This study was approved by the Institutional Animal Care and Use Committee (IACUC). Informed patient consent was not necessary for this work. We confirm that we have read the Journal’s position on issues involved in ethical publication and affirm that this work is consistent with those guidelines.

